# The potential of the *Beauveria bassiana* MHK isolate for mosquito larval control

**DOI:** 10.1101/2022.08.12.503796

**Authors:** Patil Tawidian, Qing Kang, Kristin Michel

## Abstract

The African malaria mosquito, *Anopheles gambiae* Giles (Diptera: Culicidae), and the Asian tiger mosquito, *Aedes albopictus* Skuse (Diptera: Culicidae) are of public health concern due to their ability to transmit disease-causing parasites and pathogens. Current mosquito control strategies to prevent vector-borne diseases rely mainly on the use of chemicals. However, insecticide resistance in mosquito populations necessitates alternative control measures, including biologicals such as entomopathogenic fungi. Here we report the impact of a *Beauveria bassiana* (Balsamo) Vuillemin (Hyprocreales: Cordycipitaeceae) isolate, MHK, isolated from field-collected *Ae. albopictus* larvae on mosquito survival and development. Larval infection bioassays using three *B. bassiana* conidial doses were performed on the second and third larval instars of *An. gambiae* and *Ae. albopictus* mosquitoes. Larvae were monitored daily for survival and development to pupae and adults. Our results show that *B. bassiana* MHK was more effective in killing *An. gambiae* than *Ae. albopictus* larvae. We further observed delays in development to pupae and adults in both mosquito species exposed the varying doses of *B. bassiana* as compared to the water control. In addition, larval exposure to *B. bassiana* reduced adult male and female survival in both mosquito species, further contributing to mosquito population control. Thus, this study identifies the locally isolated fungus, *B. bassiana* MHK, as a possible biological control agent of two mosquito species of public health concern, increasing the arsenal for integrated mosquito control.

## Introduction

Control of mosquito larvae through source reduction and larvicide application is an essential part of integrated mosquito management for the reduction of vector-borne diseases (Rose 2001, Floore 2006, Abdullah Shaukat et al. 2019, Fouet and Kamdem 2019). In areas where source reduction is difficult due to the inability to eliminate mosquito breeding habitats, larvicides are largely used for the control of mosquito larvae (Floore 2006). Studies have reported that repeated application of chemical larvicides has led to the development of insecticide resistance in mosquito populations (Kudom et al. 2012, Vontas et al. 2012, Wang et al. 2020). Microbial larvicides, including *Bacillus thuringiensis* var. *israelensis* [Bti] and *Bacillus sphaericus* have been used as alternatives to chemical insecticides for the control of mosquito larvae in the field (Chapman 1974, Ramírez-Lepe and Ramírez-Suero 2012). While development of resistance towards microbial larvicides are less common than chemical insecticides, studies have reported resistance to *B. sphaericus* and *B. thuringienses* in *Culex quinquefasciatus* Say (Diptera: Culicidae) larvae (Mulla et al. 2003, Wirth et al. 2004, Yuan et al. 2010). Thus, identifying novel biological control agents, including entomopathogenic fungi are crucial for the sustainable control of mosquitoes of public health concern.

The potential use of fungi as entomopathogens to control mosquito larvae and adults has been reported in mosquitoes belonging to numerous genera, including *Anopheles, Aedes*, and *Culex* (reviewed in e.g. Christophers 1952, Jenkins 1964, Roberts 1974, Castillo and Roberts 1980, Scholte et al. 2004, Tawidian et al. 2019). An example is *Metarhizium anisopliae* (Metschn.) Sorokin (Hypocreales: Clavipitaceae), an opportunistic fungal entomopathogen that has shown promise for the control of larvae and adults of the African malaria mosquito, *Anopheles gambiae* Giles (Diptera: Culicidae) in laboratory and semi-field environments (Bukhari et al. 2011, Fawrou et al. 2013, Lovett et al. 2019). Similarly, the ascomycetous fungus, *Beauveria bassiana* (Balsamo) Vuillemin (Hypocreales:Cordycipitaceae), is an entomopathogenic fungus that has a wide host range, including larvae and adults of several mosquito species (Clark et al. 1967, 1968, Miranpuri and Khachatourians 1991, Geetha and Balaraman 1999, Bukhari et al. 2010, 2011, Darbro et al. 2012, Ishii et al. 2017, Benzina et al. 2018). *B. bassiana* infects mosquito larvae by attaching to the larval body at the head and perispiracular lobes of the siphon (Clark et al. 1968, Miranpuri and Khachatourians 1991) and through ingestion of fungal spores (Miranpuri and Khachatourians 1990). Infection of mosquito larvae by *B. bassiana* causes histological changes in the larval body, including the disintegration and deformation of larval cuticle, epidermis, and adipose tissue (Benzina et al. 2018).

The overall goal of this study was to assess the entomopathogenic activity of a new isolate of *B. bassiana* towards two mosquito species of public health concern, the African malaria mosquito, *An. gambiae*, which vectors human malaria-causing parasites in sub-Saharan Africa (Craig et al. 1999), and the Asian tiger mosquito, *Aedes albopictus* Skuse (Diptera: Culicidae), which vectors the causative agents of dengue fever, chikungunya, and Zika (Gubler 2002, Girard et al. 2020). Here, we isolated *B. bassiana* MHK from *Ae. albopictus* larvae collected from Manhattan, KS, USA and assessed its impact on mosquito larval survival and development to adulthood in detail. We conducted infection bioassays by exposing second instar (L2) and third instar (L3) *An. gambiae* and *Ae. albopictus* larvae to varying doses of *B. bassiana* MHK and a commercially available *B. bassiana* isolate, GHA. Daily larval survival and development to pupae and adults were recorded. We determined that the larvicidal activity of *B. bassiana* MHK was more evident in *An. gambiae* larvae than *Ae. albopictus*. In addition, we show that larval exposure to *B. bassiana* MHK, irrespective of mosquito species, delays larval development to pupae and adults, and further reduces the survival of male and female mosquito adults. Finally, we show that *B. bassiana* MHK was as effective as the commercially available isolate, GHA, in reducing mosquito larval survival and delaying development to pupae and adults.

## Materials and methods

### Isolation of fungi from field-collected *Ae. albopictus* larvae

Mosquito-associated fungi were isolated from fourth instar *Ae. albopictus* larvae collected in Manhattan, KS during 2018. The larval breeding sites consisted of human-made mosquito oviposition cups lined with heavyweight seed germination paper (Anchor Paper Co., MN, USA). Individual larvae were surface washed six times with 1% phosphate-buffered saline and ground whole in 50 μL sterile water. To prevent bacterial growth, each larval suspension was spread on potato dextrose agar (PDA) plates (BD Difco™, NJ, USA) supplied with 100 mg/mL ampicillin (Sigma-Aldrich, MO, USA) and 50 mg/mL chloramphenicol (Sigma-Aldrich, MO, USA). PDA plates were then incubated at room temperature in the dark for five days. Thereafter, unique fungal morphotypes were identified and transferred to new PDA plates and incubated at room temperature for 14 days. Preliminary identification of the locally isolated *B. bassiana*, hereafter referred to as MHK isolate, was performed by analyzing the morphological characteristics under an optical microscope (400x). In addition, a commercialized isolate of *B. bassiana*, isolate GHA, the active ingredient in Aprehend^®^ (ConidioTec LLC, Centre Hall, PA, USA), PA, USA), was maintained on PDA media following the same incubation procedure described above.

### Molecular identification of *B. bassiana* isolates

Total genomic DNA was isolated from mycelia and conidia of the *B. bassiana* isolates using the Dneasy PowerSoil kit (MoBio Laboratory, Carlsbad, CA, USA) according to the manufacturer’s instructions, with the minor modification of increasing the time of centrifugation at 11,000 rpm to 5 minutes to ensure the removal of residual ethanol from the MB spin column. *B. bassiana* MHK and GHA isolates were further characterized through the amplification of the fungal Internal Transcribed Spacer marker using forward primer ITS1f (5’-CTTGGTCATTTAGAGGAAGTAA-3’) (Gardes and Bruns 1993) and reverse primer ITS4 (5’-TCCTCCGCTTATTGATATGC-3’) (White et al. 1990) according to published protocols (Ihrmark et al. 2012) with minor modification. PCR with 20 ng of template DNA included an initial denaturation step for 5 min at 94°C and 35 cycles of denaturing, annealing, and extension at 94°C for 30 s, 58°C for 30 s, and 72°C for 30 sec, followed by a final 72°C extension step for 7 min. The PCR also included a negative control consisting of certified nuclease-free water. Successful PCR amplification was determined by visualizing 5 μL of the PCR products on 1% agarose gels. The remaining 45 μL of PCR products were purified using a Qiaquick® PCR purification kit (Qiagen, Germany). The purified PCR amplicons were sequenced by Sanger sequencing at GeneWiz. The ITS region sequence of *B. bassiana* MHK isolate was submitted to GenBank, and assigned Accession # OP186482.

### Mosquito rearing

*An. gambiae* (G3 strain) and *Ae. albopictus* (10^th^ to 20^th^ generations collected in Missouri and colonized by the Illinois Natural History Survey) mosquitoes were reared at 27°C and 80% relative humidity using a 12L:12D photoperiod cycle. *An. gambiae* larvae were reared as described previously (An et al. 2011), with freshly hatched larvae being fed on 20 g/L baker’s yeast (Active Dry Yeast, Fleischmann’s) for 48 hours followed by a mixture of 6.6 g/L baker’s yeast and 13.4 g/L fish food (TetraMin® Tropical Flakes, Tetra). *Ae. albopictus* eggs were hatched as described previously (Zheng et al. 2015) in water containing 0.36 g/L of CM0001 Nutrient Broth (Oxoid, Hampshire, England). Hatched larvae were fed daily on a mixture of baker’s yeast and fish food (TetraMin® Tropical Flakes, Tetra). Adult mosquitoes of both species were provided with a sugar solution containing 8% fructose (Emprove® Chemicals, Sigma, MO, USA) *ad libitum*.

### Conidial suspension and viability

*B. bassiana* isolates MHK and GHA were grown separately in Petri dishes (150 mm x 15 mm) containing PDA media and incubated at room temperature for 14 days. For each isolate of *B. bassiana*, conidia were scraped from 14 petri dishes containing PDA media colonized by fungal mycelia using sterile water and T-spreaders. Conidial suspensions were filtered through four layers of sterile cheesecloth and transferred to sterile conical tubes. Conidial concentrations were determined using a hemocytometer and adjusted to 2.5×10^7^ conidia/mL, 1.25×10^8^ conidia/mL, and 2.5×10^8^ conidia/mL, hereafter referred to as low, medium, and high doses. New conidial suspensions were prepared for each bioassay and used immediately after suspension in water. Prior to each bioassay, the viability of conidia was determined by placing 100 μL of conidial suspensions adjusted at 10^5^ conidia/mL on PDA media. We then observed under optical microscopy (400x) whether 300 conidia had germinated 18 h after inoculation. Conidia with germ tubes twice their diameter were considered viable (Francisco et al. 2006).

### Bioassays

To assess the impact of larval exposure to *B. bassiana* conidia on mosquito survival and development, we used infection bioassays in a fully factorial arrangement that included 32 individual combinations of target population and treatments. The experiment was designed to test the impact of the following four variables on infection outcome: (i) mosquito species, using *Ae. albopictus* and *An. gambiae* larvae; (ii) larval age/stage, using L2 and L3 instars; (iii) dose, using three conidial concentrations and a water-only control, and (iv) fungal isolate, comparing *B. bassiana* MHK to the GHA isolate. All infection bioassays were performed in 24-well plates. Each well contained 1 mL of water including conidia at the low, medium, and high dose or water as control, respectively. Mosquito larvae, either as L2 or L3 were then added immediately to each well, so that each well only contained one larva to prevent cannibalism. Larvae were fed daily by adding 20 μl of food slurry (6.6 g/L baker’s yeast, 13.6 g/L fish food in Milli-Q® purified water) to each well. Each of the 32 combinations were performed in three 24-well plates (n=72 larvae). To account for batch effects, each combination was repeated thrice at separate times, yielding three trials, with a total n=216 mosquito larva per combination. Each larva was monitored daily for survival, death, and pupation. Pupae were collected daily, washed with sterile water, separated based on pupation date and treatment, and placed into pint-sized emergence cups (Neptune Paper Inc, Fort Lee, NJ, USA) covered with tulle netting (Jo-Ann Fabric and Crafts, Hudson, OH, USA). Emergence cups were monitored daily for pupal survival, death, and eclosion of female and male adult. Adults were fed ad libitum using cotton balls, which were soaked in 8 % fructose and placed on top of the emergence cups. Male and female mosquitoes were monitored daily for death for ten days post eclosion.

### Statistical Analysis

Statistical analyses were executed using the MIXED, LOGISTIC, PHREG and LIFETEST procedures in SAS/STAT® software, version 9.4 (SAS Institute Inc., Cary, NC). We analyzed the survival proportions from L1 to pupa and L1 to sex-specific adults (Supp Tables S1-S4) using conditional logistic regression model with fungal dose as a fixed effect and trial forming strata. When survival in a pairwise comparison was rare, the exact inference was employed in hypothesis testing and estimation of odds ratios (OR). The survival proportion of each fungal dose was estimated by averaging the survival proportion across trials. In addition, we analyzed larval development time as measured by time to pupation and time to eclosion (L1 being Day 0) using a linear mixed model with fungal dose and trial as fixed effects. Residual variance was allowed to be heterogeneous with respect to fungal dose. The least squares means (LSMs) and their standard errors (SEMs) were reported for each fungal dose.

Sex-specific adult survival post-eclosion (Eclosion being Day 0) was analyzed using Cox’s proportional hazard model with fungal dose being the fixed effects and trial forming strata. The hazard ratios (HRs) were used to compare fugal doses. Adult survival curves up to ten days post-eclosion were obtained via the Kaplan-Meier estimator across trials. Pairwise comparisons among dose levels were performed based on the two-sided test for non-zero difference in means, non-one ratio in odds, or non-one ratio in hazard rate. No multiplicity adjustment was applied. In addition, statistical analyses of adult survival proportions from some fungal doses were excluded due to high larval mortality.

## Results

### Morphological and sequence characterization of the MHK isolate of *B. bassiana*

*B. bassiana* MHK was isolated from *Ae. albopictus* mosquito larvae collected from Manhattan, KS, USA. *B. bassiana* MHK fungal colony morphology was compared to the commercial *B. bassiana* GHA isolate (Fig. 1). Seven days after plate inoculation, colony diameters were 22 cm and 20 cm for MHK and GHA isolates, respectively. On PDA, both *B. bassiana* isolates had white mycelial growth on the surface and a reverse pale orange to yellow growth (Fig. 1A and B). While both fungal isolates had filamentous colony margins, the margins were more distinct in *B. bassiana* GHA compared to MHK (Fig. 1A and B). Conidia of both *B. bassiana* isolates ranged between 2.00-2.37 μm and were hyaline and globose in shape (Fig. 1C and D).

**Fig. 1.**
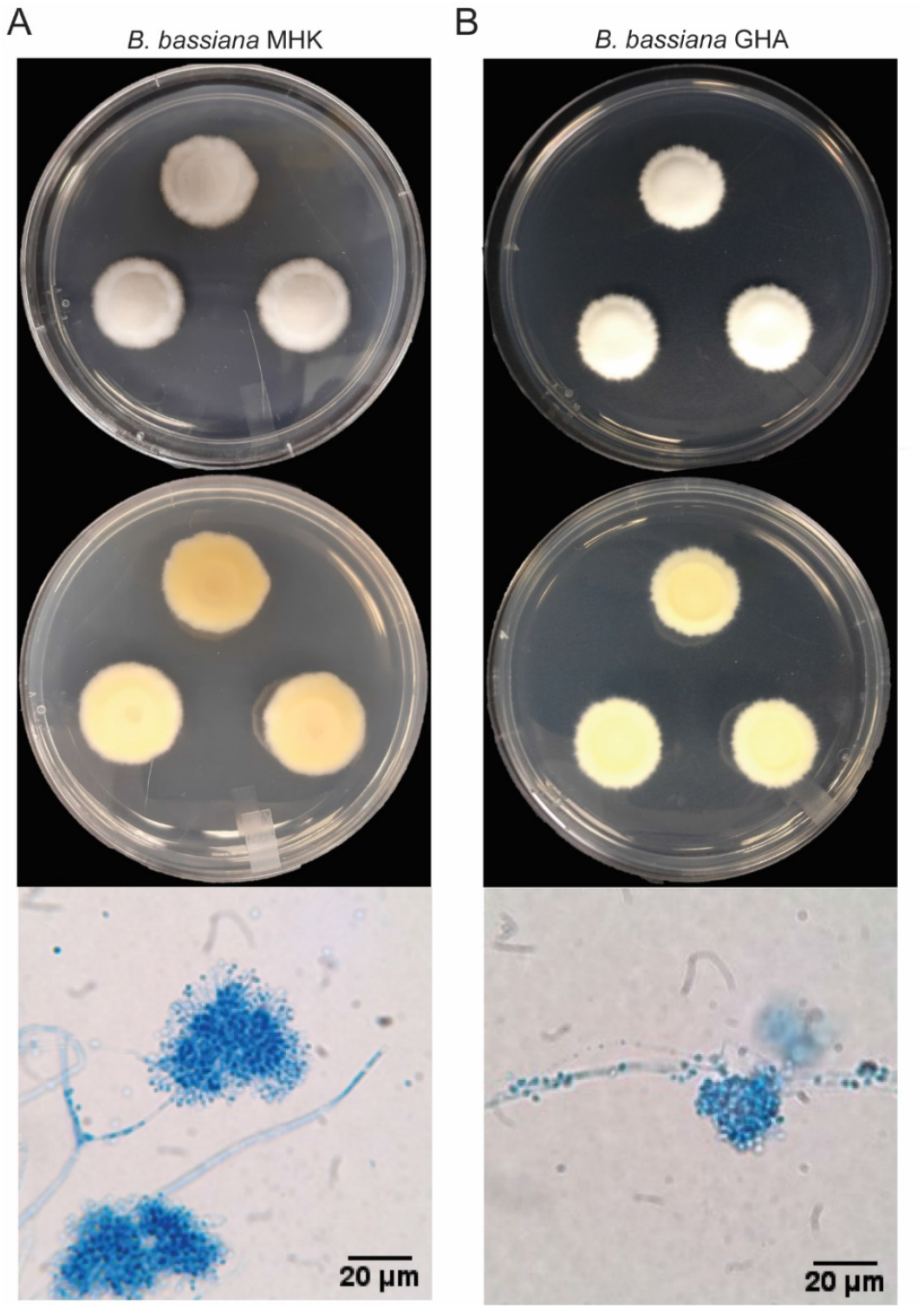
*B. bassiana* MHK and GHA mycelial growth characteristics and conidial morphology. (**A**) Top and bottom view of *B. bassiana* MHK mycelial growth on potato dextrose agar (PDA) media and a microscopic overview of the conidiophore and fungal conidia; (**B**) Top and bottom view of *B. bassiana* GHA mycelial growth on PDA media and a microscopic overview of the conidiophore and fungal conidia.

In addition to morphology-based characterization, the ITS1, 5.8S, and ITS2 regions of the MHK isolate were sequenced. Nucleotide sequence comparison to the nonredundant GenBank database revealed 99-100% sequence identity to known *B. bassiana* isolates, confirming the fungal species identification based on morphology.

### *B. bassiana* MHK and GHA reduced larval survival in both mosquito species

To assess the impact of *B. bassiana* isolates on mosquito larval survival, we plotted larval survival proportions and compared the odds of larval survival (L1 to pupa) across fungal doses and water control (Table 1, Fig. 2). In the absence of conidia 93.1 – 100 % of *An. gambiae* larvae and 95.8 – 99.5 % of *Ae. albopictus* larvae developed to the pupal stage, indicating that our experimental setup did not impair progression of larval to pupal stages.

**Table 1.**
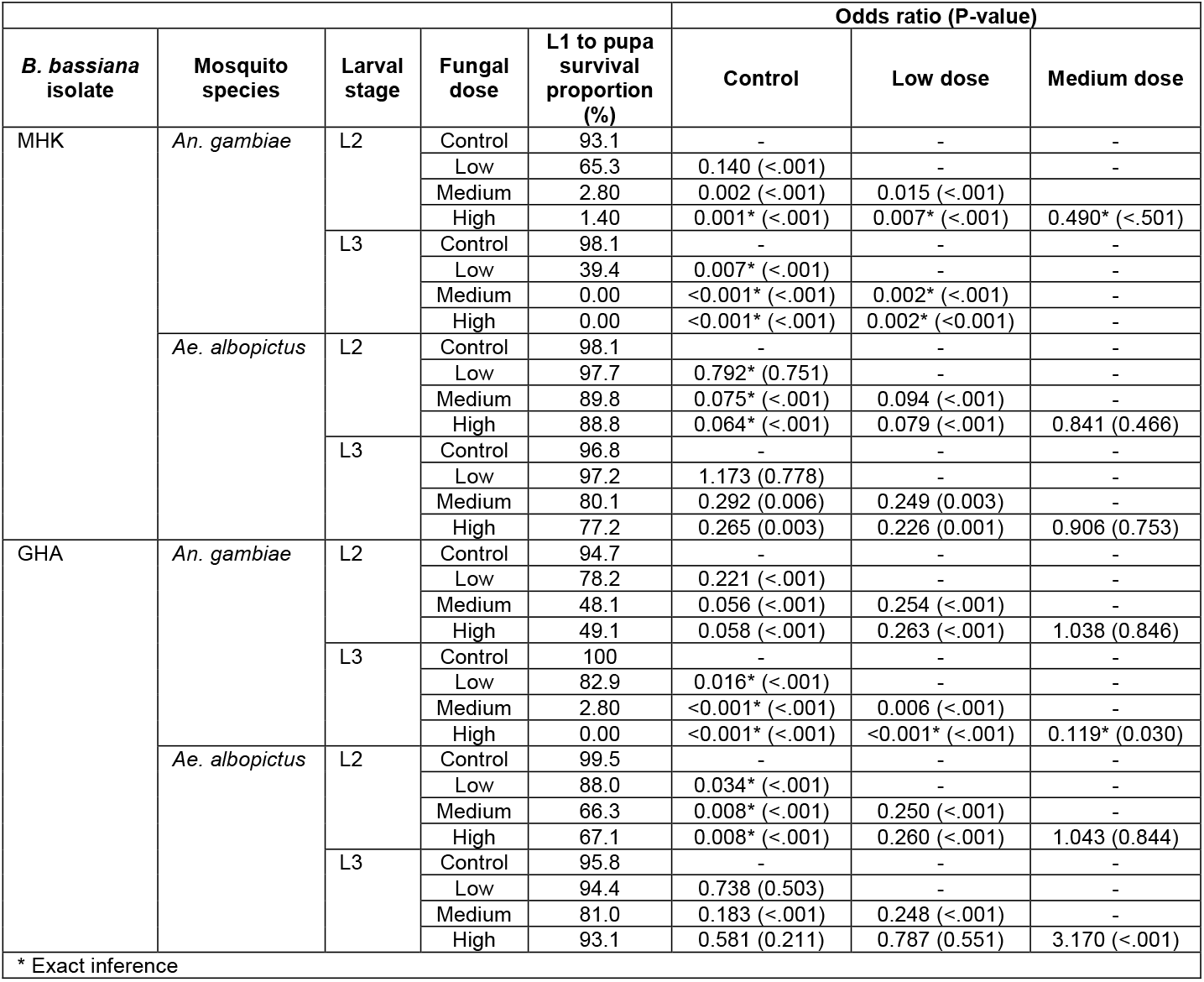
Odds ratios of *An. gambiae* and *Ae. albopictus* L1 to pupa survival exposed to *B. bassiana* MHK and GHA as L2 and L3 larvae.

**Fig. 2.**
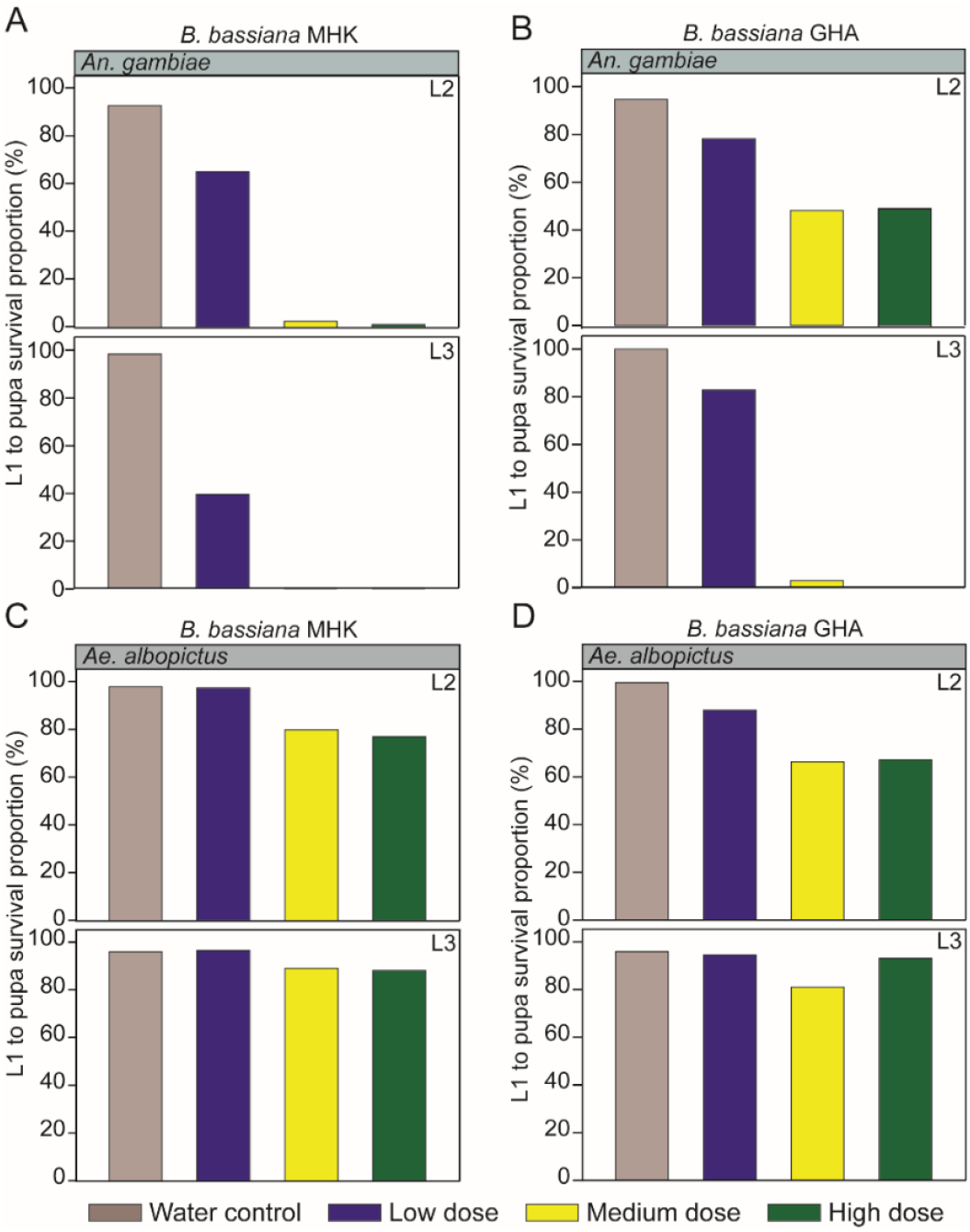
Survival proportions (%) of L1 larvae to pupae for larvae exposed as L2 and L3 instars to *B. bassiana*. Larval exposure stage to *B. bassiana* is indicated at the top right of each bar graph, while the keys at the bottom of the plots depict the fungal doses and water control color coded differently. (**A**) L1 to pupa survival proportion of *An. gambiae* larvae exposed to *B. bassiana* MHK; (**B**) L1 to pupa survival proportion of *An. gambiae* exposed to *B. bassiana* GHA; **(C)** L1 to pupa survival proportion of *Ae. albopictus* larvae exposed to *B. bassiana* MHK; and (**D**) *Ae. albopictus* larvae exposed to *B. bassiana* GHA.

Exposure to *B. bassiana* MHK conidia significantly reduced the odds of *An. gambiae* larval survival compared to the water control, irrespective of mosquito stage at exposure (Table 1). Nearly all larvae exposed as L2 and L3 to the medium and high fungal doses died (0% survival of larvae exposed as L3 to medium or high dose; 1.4 % and 2.8% survival of larvae exposed as L2 to medium and high dose, respectively, Fig. 2A). However, survival was less impaired when larvae were exposed to the lowest fungal dose (39.4 and 65.3% survival of larvae exposed as L3 and L2, respectively, Fig. 2A). The pattern of *An. gambiae* larval survival exposed as L3 to *B. bassiana* GHA remained overall similar to the pattern observed with the MHK isolate. However, survival proportions of larvae exposed to medium and high doses of the GHA isolate as L2 were 17- and 35-fold higher than those exposed to the same doses of the MHK isolate (Table 1, Fig. 2B).

The impact of *B. bassiana* on *Ae. albopictus* larval survival was overall much smaller compared to the impact on *An. gambiae* larvae, irrespective of the fungal isolate (e.g., 77.2% and 67.1% survival of larvae exposed as L2 to high dose of MHK and GHA isolate, respectively Table 1, Fig. 2C and D). Nevertheless, the odds of *Ae. albopictus* larval survival was decreased significantly when exposed to the medium and high fungal doses of the *B. bassiana* MHK isolate (Table 1). Survival percentages of *Ae. albopictus* larvae exposed to the medium and high MHK doses were reduced to 77.2 and 89.8% when exposed as L3, as compared to >96% survival for larvae exposed to the low dose and water control (Fig. 2C). In contrast, all doses of *B. bassiana* GHA significantly reduced larval survival in *Ae. albopictus* larvae exposed as L2, but only the medium dose reduced larval survival (to 81%) when exposed as L3, as compared to the water control (Table 1, Fig. 2D).

Therefore, the MHK isolate overall showed similar efficacy in larval killing as the commercial GHA isolate, irrespective of mosquito species and larval stage at exposure. However, *B. bassiana* was more efficacious in killing *An. gambiae* than *Ae. albopictus*, irrespective of fungal isolate and larval stage at exposure.

### The *B. bassiana* MHK isolate reduced pupal survival in *Ae. albopictus*

To assess whether larval exposure to *B. bassiana* impacted mosquito survival at the pupal stage, we plotted L1 to adult survival proportions and compared the percent survival to adulthood with the percent survival to pupae across the different doses and water control (Tables 1 and 2, Figs. 2 and 3). In the absence of conidia, 78.7 – 89.8 % of *An. gambiae* larvae and 88.4 – 94.0 % of *Ae. albopictus* larvae developed to the adult stage, revealing that 9.4 – 15.4 % of *An. gambiae* and 5.2 – 8.6% of *Ae. albopictus* died in the pupal stage.

**Table 2.**
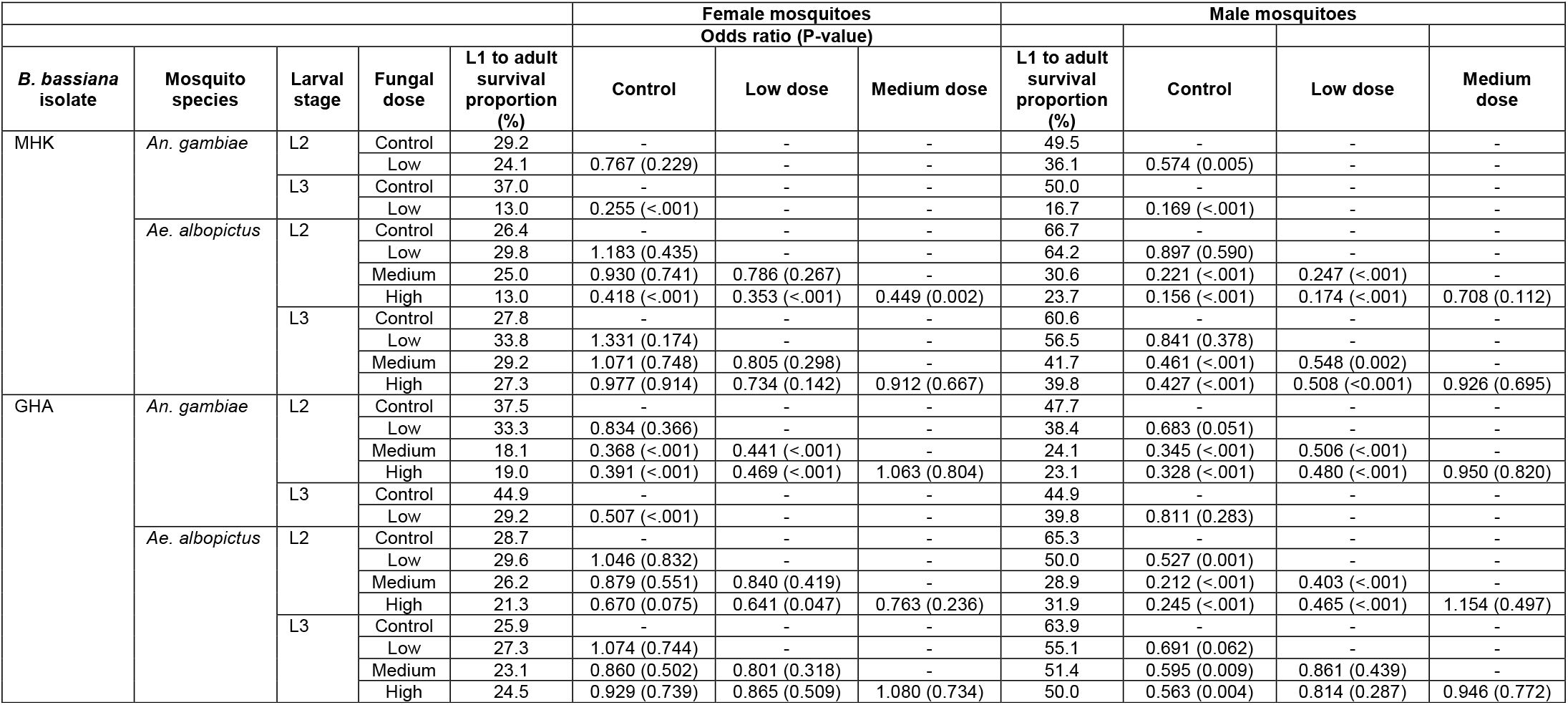
Odds ratios of *An. gambiae* and *Ae. albopictus* L1 to adult survival exposed to *B. bassiana* MHK and GHA as L2 and L3 larvae.

**Fig. 3.**
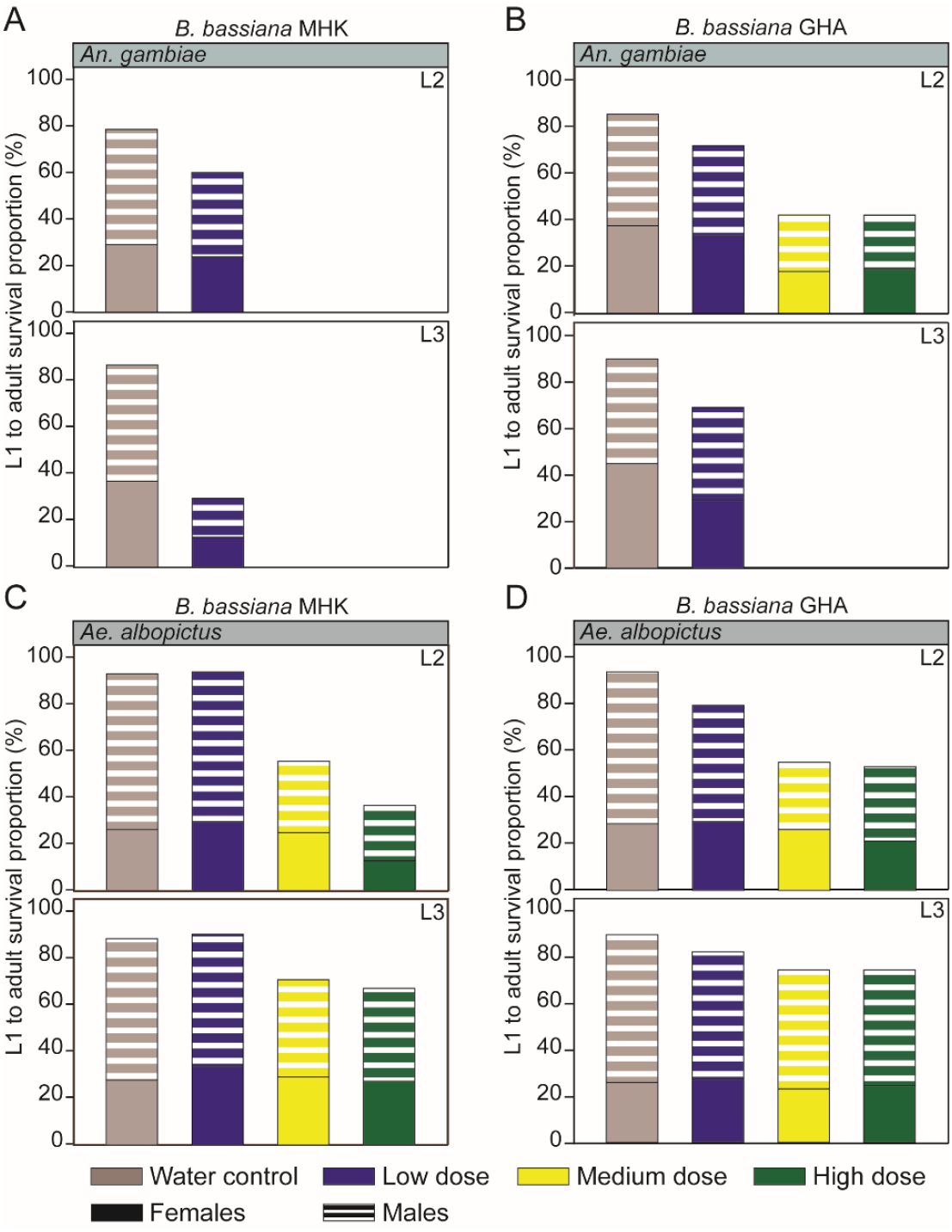
Survival proportions (%) of L1 larvae to adults for larvae exposed as L2 and L3 instars to *B. bassiana*. Larval exposure stage to *B. bassiana* is indicated at the top right of each bar graph. Solid bars indicate adult female mosquito survival proportions, while striped bars depict those of adult male mosquitoes. L1 to adult *An. gambiae* survival proportions for larvae exposed to *B. bassiana* MHK **(A)** and GHA **(B)** isolates; L1 to adult *Ae. albopictus* survival proportions for larvae exposed to *B. bassiana* MHK **(C)** and **(D)** GHA isolates.

The impact of *B. bassiana* on *An. gambiae* pupal survival was overall small, as the majority of mosquitoes had died at the larval stage after fungal exposure. The few *An. gambiae* larvae that developed to the pupal stage died as pupae. We observed no pupal survival of *An. gambiae* exposed as either L2 or L3 larvae to medium and high doses of *B. bassiana* MHK and exposed as L3 larvae to medium and high doses of *B. bassiana* GHA (Fig. 3A and B). In the one instance where a substantial number of *An. gambiae* larvae survived the exposure to *B. bassiana* (medium or high dose GHA conidia, exposure at L2), no additional pupal death was observed (Tables 1 and 2, Figs. 2B and 3B).

In contrast, while medium and high doses of both isolates of *B. bassiana* had limited impact on *Ae. albopictus* larval survival, the same treatments caused substantial pupal mortality (Table 2, Fig. 3C and D). Overall, high dose exposure to *B. bassiana* killed 14.0 – 40.5 % of pupae, dependent on fungal isolate and larval exposure stage (Tables 1 and 2). Lowering the dose reduced pupal death, as exposure to a medium dose of conidia killed 6.5 – 24.5 % of pupae, dependent on fungal isolate and larval exposure stage. Interestingly, the MHK isolate caused more pupal mortality than the GHA isolate, irrespective of larval exposure stage and conidial dose. This was most pronounced after exposure of L2 larvae to high dose conidia, where the MHK isolate reduced pupal survival by 40.5 %, while the GHA isolate reduced pupal survival by only 14 % (Tables 1 and 2).

Taken together, these results show that *B. bassiana* isolates had minimal impact on *An. gambiae* pupal survival. Both *B. bassiana* isolates reduced significantly *Ae. albopictus* pupal survival, and the MHK isolate reduced pupal survival more strongly than the GHA isolate.

### Sex-specific reduction in the survival of *Ae. albopictus* mosquito larvae and pupae exposed to *B. bassiana*

To assess whether larval exposure to *B. bassiana* impacted mosquito survival in a sex-specific manner, we plotted L1 to adult survival proportions and compared the odds ratio of survival to adults between treatments separately for males and females (Table 2, Fig. 3). In the absence of conidia, 24.2 – 44.9 % of *An. gambiae* larvae developed to adult females, while 41.1 – 50.0 % of *An. gambiae* larvae developed to adult males. For *Ae. albopictus*, in the absence of conidia, 25.9 – 28.7 % of larvae developed to adult females, while 60.6 – 66.6 % of larvae developed to adult males.

Exposure to *B. bassiana* impacted *An. gambiae* male and female larval/pupal survival, irrespective of fungal isolate and exposure larval stage (Table 2, Fig. 3A and B). While *B. bassiana* strongly impacted overall survival to adulthood of *An. gambiae*, differential impact on sex-specific larval/pupal survival was only observed in two fungal treatments (Table 2). First, the MHK isolate only reduced significantly the odds of survival to adult males and not females, when L2 larvae were exposed to a low dose of conidia as compared to the water control (OR=0.574, *P*=0.005 for males; OR=0.767, *P*=0.229 for females). Second, the GHA isolate only affected the odds of survival to adult females and not males when L3 larvae were exposed to a low dose as compared to the water control (OR=0.811, *P*=0.283 for males; OR=0.507, *P*<0.001 for females).

In contrast, in all *B. bassiana* treatments that reduced survival of *Ae. albopictus* larvae to adults, this reduction was solely due to male larval/pupal death (Table 2, Fig. 3C and D). This differential effect of killing male larvae/pupae while leaving female larvae and pupae unharmed was observed for both *B. bassiana* isolates, and unaffected by larval exposure stage. For example, exposure to GHA conidia at the L2 stage killed an additional 33.4% of male larvae/pupae, as compared to only an additional 7.4% of female larvae/pupae as compared to water controls (OR=0.245, *P*<0.001 for males; OR=0.670, *P*=0.075 for females Table 2, Fig. 3D). The only treatment that reduced survival of both *Ae. albopictus* male and female larvae/pupae was the exposure of L2 larvae to the high dose of MHK conidia (OR=0.156, *P*<0.001 for males; OR=0.418, *P*<0.001 for females, Table 2). However, the treatment impacted males more strongly than females, killing an additional 43% of male larvae/pupae, while only killing an additional 13.4% of female larvae/pupae as compared to the water controls (Fig. 3C). In summary, our data show limited evidence that *B. bassiana* kills *An. gambiae* during development in a sex-specific manner. However, *B. bassiana* caused a strong sex-specific impact on survival of *Ae. albopictus* after larval exposure, killing mostly male larvae/pupae. In addition, the MHK isolate, dependent on fungal dose and exposure larval stage, also killed female *Ae. albopictus* during development.

### Exposure to *B. bassiana* delayed development in both mosquito species

To determine whether, in addition to killing immature mosquito stages, *B. bassiana* also delayed larval and pupal development, we assessed time to pupation and time to eclosion (Table 3, Fig. 4). In the absence of conidia, *An. gambiae* larvae developed to pupae in 8.1 – 9.1 days and adults eclosed 1.1 – 1.2 days later, while *Ae. albopictus* larvae developed to pupae in 6.4 – 6.7 days and eclosed after an additional 1.9 – 2.1 days.

**Table 3.**
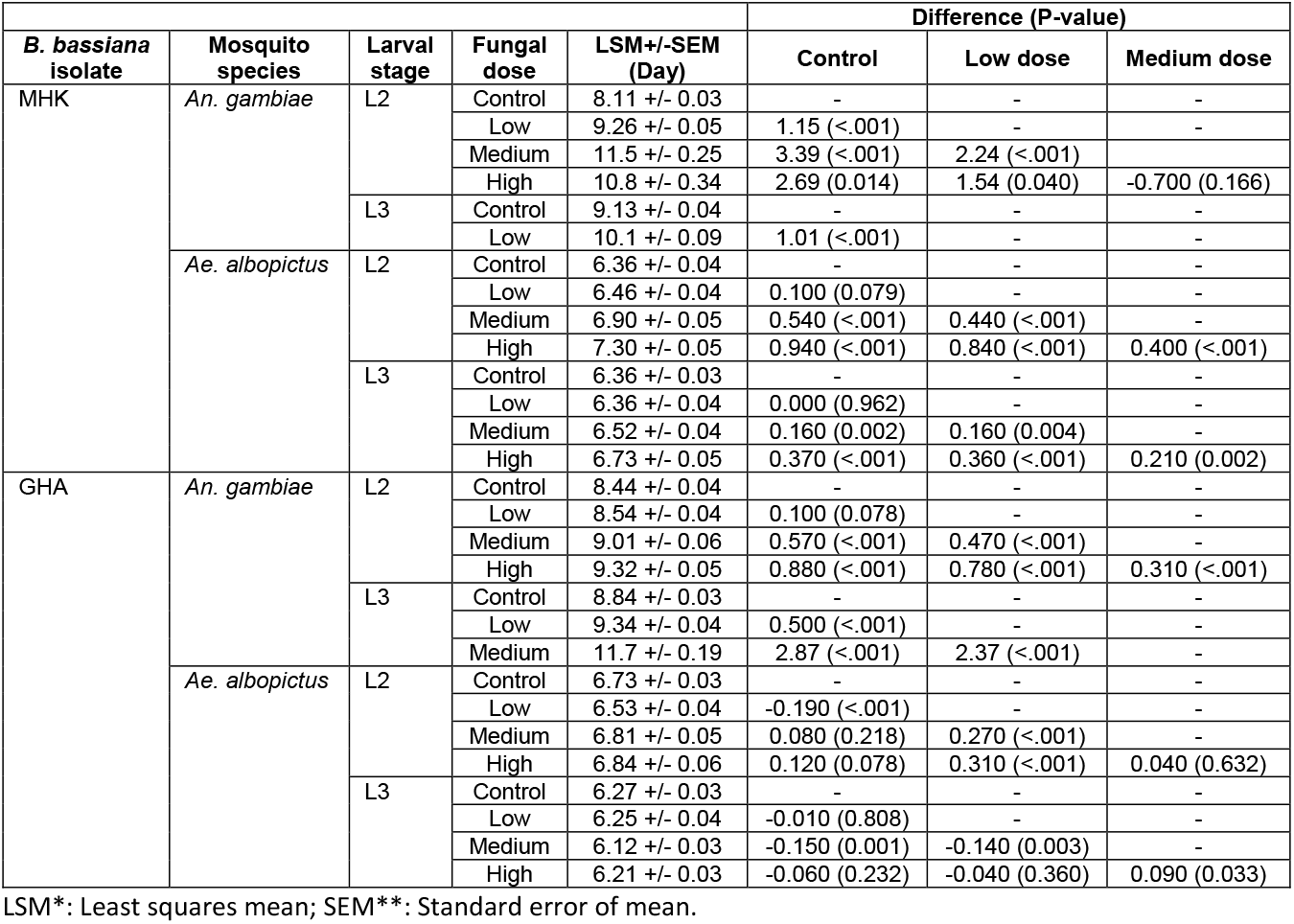
Time to pupation of *An. gambiae* and *Ae. albopictus* L2 and L3 larvae exposed to *B. bassiana* MHK and GHA.

**Fig. 4.**
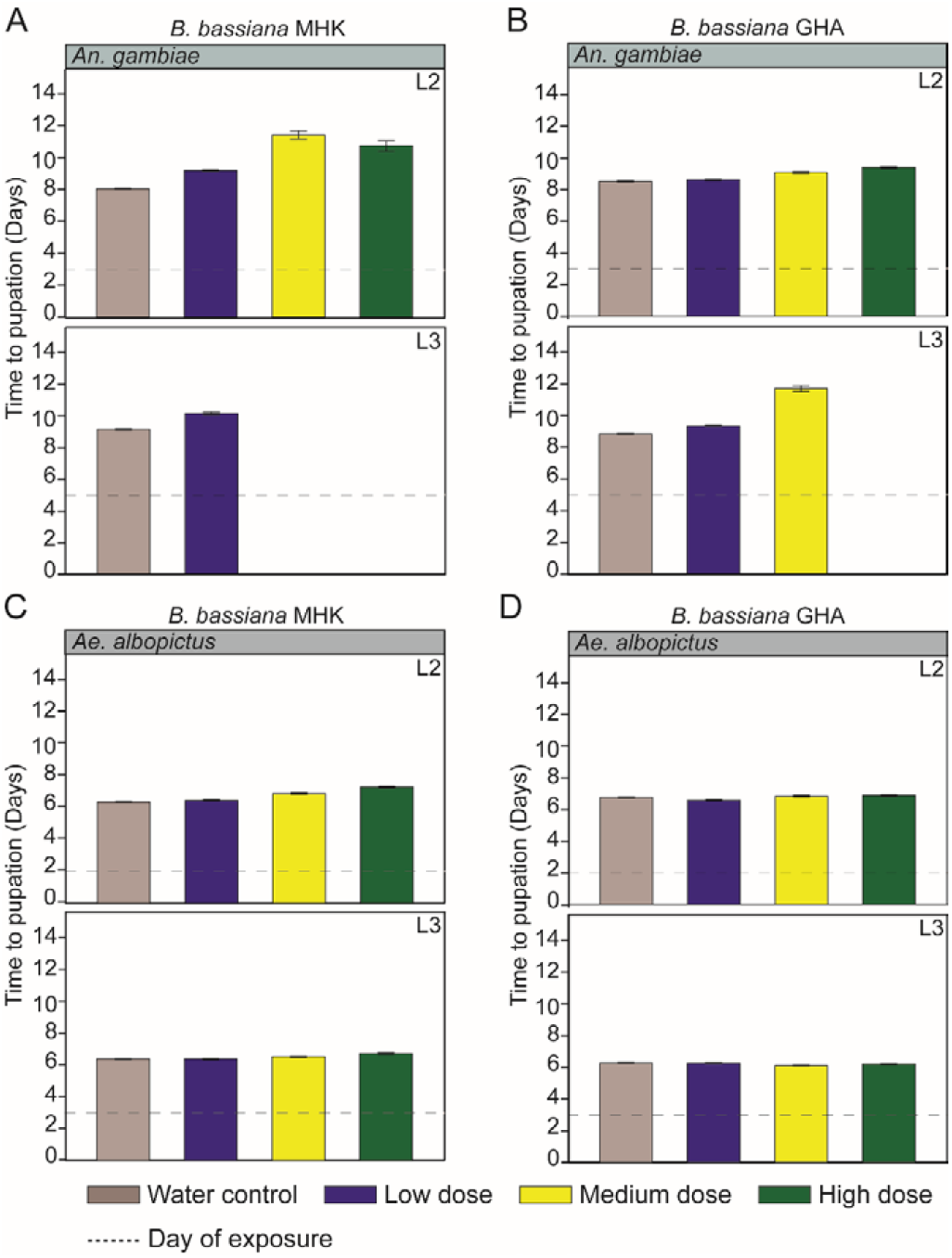
Mean time to pupation in days (L1 being Day 0) of L2 and L3 larvae exposed to *B. bassiana*. Larval exposure stage to *B. bassiana* is indicated at the top right of each bar graph. Dotted line indicates the day each larval stage was exposed to *B. bassiana*. The error bar indicates the standard error of mean time to pupation. (**A**) Mean time to pupation of *An. gambiae* larvae exposed to *B. bassiana* MHK; (**B**) Mean time to pupation of *An. gambiae* larvae exposed to *B. bassiana* GHA; (**C**) Mean time to pupation of *Ae. albopictus* larvae exposed to *B. bassiana* MHK; (**D**) Mean time to pupation of *Ae. albopictus* larvae exposed to *B. bassiana* GHA.

Exposure of *An. gambiae* larvae to *B. bassiana* delayed time to pupation, irrespective of fungal isolate and larval exposure stage (Table 3 and Supp Table S5, Fig. 4A and B and Supp Fig. S1A and B). This impact was dose-dependent, and most strongly observed for medium and high doses of the MHK isolate at L2 exposure, and for the medium dose of the GHA isolate at L3 exposure, leading to a maximum 2.8 – 3.4-day delay in time to pupation as compared to the water controls (Table 3, Fig. 4A and B). At the lowest dose, where the impact on survival was the smallest, time to pupation was reduced by at least one day when larvae were exposed to the MHK isolate (Table 3, Fig. 4A). Overall, we did not observe an additional delay in development at the pupal stage after *B. bassiana* exposure, as the maximum additional time spent as pupae was less than 0.25 days as compared to the water controls (Table 3 and Supp Table S5). The only exception was the treatment with medium dose of MHK of *An. gambiae* L2 larvae, which extended the time spent in the pupal stage by nearly one full day, though this delay was only observed for three individual pupae (Table 3 and Supp Tables S1 and S5).

In contrast, delay in *Ae. albopictus* time to pupation was only observed when larvae were exposed to *B. bassiana* MHK and not GHA (Table 3, Fig. 4C and D). While statistically significant delays in time to pupation were recorded in *Ae. albopictus* larvae exposed as L2 and L3 to *B. bassiana* MHK medium and high doses, these delays remained less than one day compared to the water control (Table 3, Fig. 4B). No additional delay in development time to eclosion was observed at the pupal stage of *Ae. albopictus*, where the largest additional time spent in the pupal stage was less than 0.23 days as compared to water controls (Table 3 and Supp Table S5).

Together, our results show that the *B. bassiana* MHK isolate significantly delayed mosquito development time, irrespective of larval exposure stage and mosquito species. However, the MHK isolate caused a larger delay in time to pupation as the GHA isolate. In addition, developmental delay was more strongly observed in *An. gambiae* larvae, and caused by a delay in time to pupation rather than time spent at the pupal stage.

### Larval exposure to B. bassiana reduced mosquito survival post-eclosion

To determine the impact of larval exposure to *B. bassiana* on male and female adult survival, sex-specific survival of adult *An. gambiae* and *Ae. albopictus* mosquitoes was recorded ten days post-eclosion.

Exposure to *B. bassiana* at the larval stage reduced *An. gambiae* post eclosion survival for both male and female adults, and was influenced by fungal isolate, exposure stage, and adult sex. The largest reduction was seen after exposure to *B. bassiana* GHA, where adult male and female *An. gambiae* exposed at the L3 stage to low dose conidia were nearly three and four times more likely to die as compared to the water control, respectively (HR=2.72, *P=0.012* in males; HR=3.78, *P=0.003* in females, Table 4, Fig. 5A and B). In addition, *B. bassiana* GHA reduced survival of female, but not male adults when exposed as L2 to the highest conidial dose (HR=1.82, *P=0.165* in males; HR=2.54, *P=0.043* in females, Table 4, Fig. 5A and B). A similar reduction in adult *An. gambiae* survival was observed after exposure to the MHK isolate, but was strongly dependent on the larval exposure stage. Low dose conidia significantly reduced *An. gambiae* male adult survival, but only when exposure occurred at the L2 stage (HR= 2.42, *P=0.031* for L2; HR= 0.99, *P=0.980* for L3, Table 4, Fig. 5C). The opposite was observed for female *An. gambiae*, where low dose conidia significantly reduced female adult survival as compared to water control only when exposure occurred at the L3 stage (HR= 1.61, *P=0.465* for L2; HR=7.23, *P=0.001* for L3, Table 4, Fig. 5D). *B. bassiana* impacted *Ae. albopictus* post eclosion survival only after exposure to the highest conidial dose at the L2 stage, regardless of fungal isolate and adult sex (Table 4, Fig. 6). *Ae. albopictus* males and females exposed as L2 larvae to the high fungal dose of MHK conidia were seven and twelve times more likely to die as compared to the water control, respectively (HR=7.52, *P<0.001* in males; HR=12.2, *P=0.001* in females, Table 4, Fig. 6A and B). Similar patterns of mortality were observed in *Ae. albopictus* adults exposed as L2 larvae to the high fungal dose of GHA. Male and female adults were two and six times more likely to die as compared to the water control, respectively (HR=2.48, *P=0.036* in males; HR=6.05, *P=0.029* in females) (Table 4, Fig. 6C and D). No fungal treatment reduced adult survival of *Ae. albopictus* when exposure occurred at the L3 stage (e.g., HR=1.86, *P=0.197* in males; HR=1.76, *P=0.316* in females, MHK high dose) (Table 4).

**Table 4.**
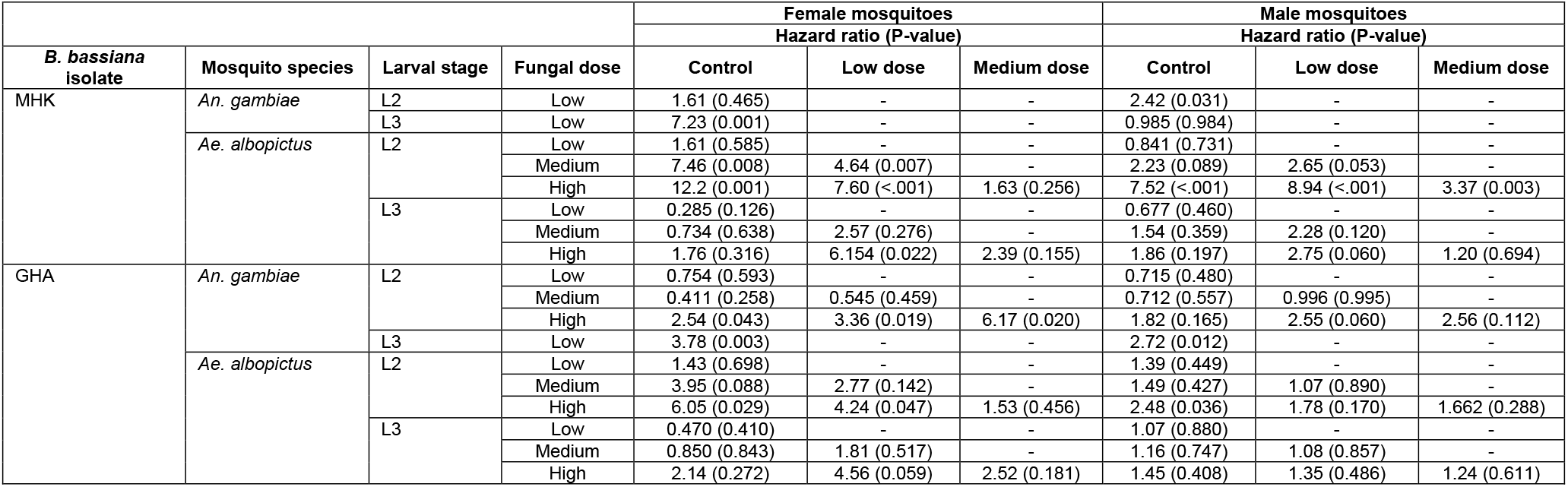
Hazard ratio of male and female *An. gambiae* and *Ae. albopictus* adults up to ten days post-eclosion exposed as L2 and L3 larvae to *B. bassiana* MHK and GHA.

**Fig. 5.**
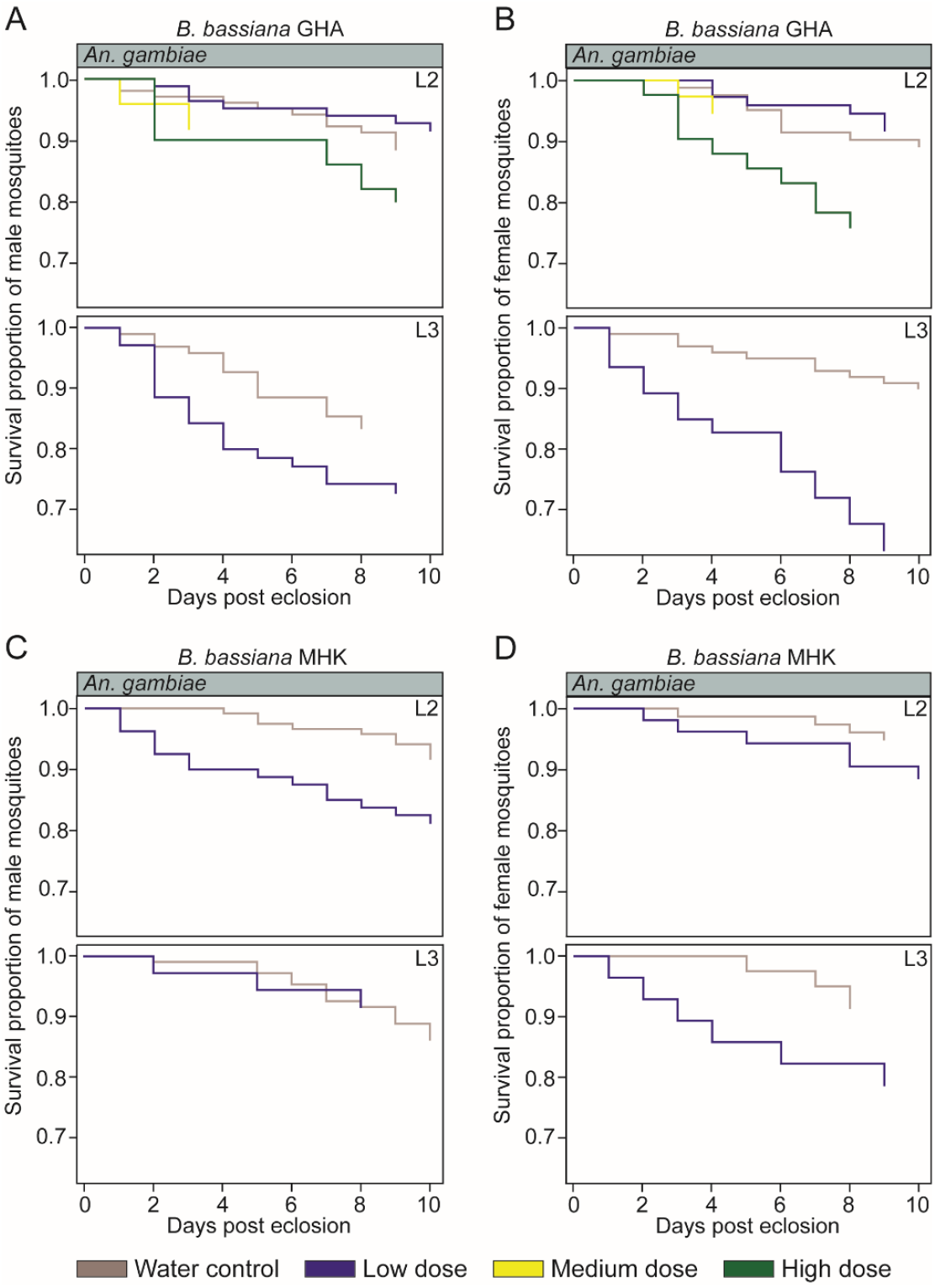
Survival proportions up to ten days post-eclosion of *An. gambiae* male and female mosquito adults exposed as larvae to *B. bassiana*. Larval exposure stage to *B. bassiana* is indicated at the top right of each bar graph. Survival proportion of *An. gambiae* male adults **(A)** and female adults **(B)** when exposed as larvae to *B. bassiana* GHA; Survival proportion of *An. gambiae* male adults **(C)** and female adults **(D)** when exposed as larvae to *B. bassiana* MHK.

**Fig. 6.**
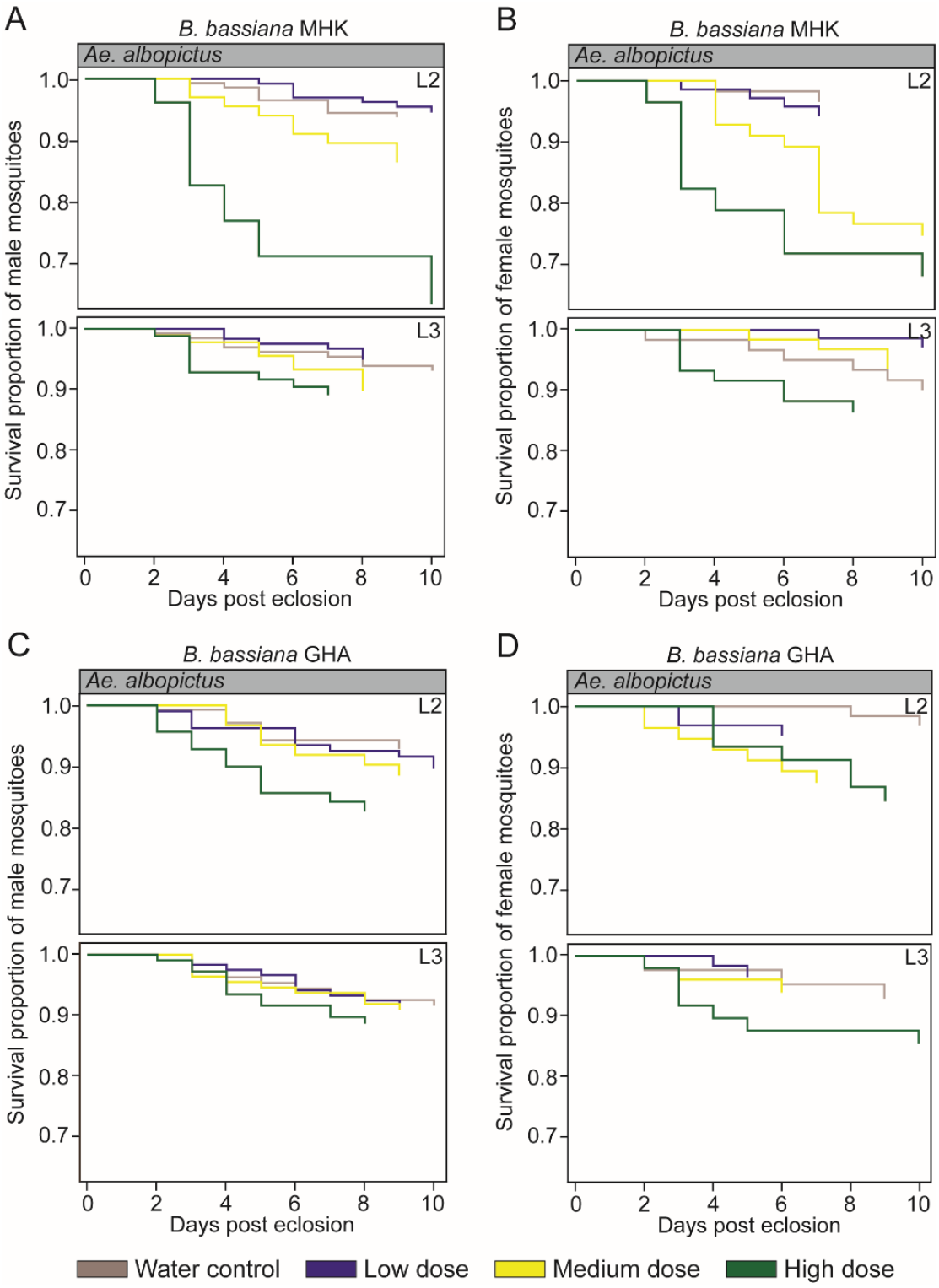
Survival proportions ten days post-eclosion of *Ae. albopictus* male and female mosquito adults exposed as larvae to *B. bassiana*. Larval exposure stage to *B. bassiana* is indicated at the top right of each bar graph. Survival proportion of *Ae. albopictus* male adults **(A)** and female adults **(B)** when exposed as larvae to *B. bassiana* MHK; Survival proportion of *Ae. albopictus* male adults **(C)** and female adults **(D)** when exposed as larvae to *B. bassiana* GHA.

In summary, these data revealed that an exposure to *B. bassiana* at the larval stage impacted post eclosion survival of both mosquito species. The effect on *Ae. albopictus* was largely limited to exposure of L2 larvae to the highest dose of conidia and not influenced by fungal isolate, or adult sex. In contrast, the reduction of *An. gambiae* post eclosion survival was measurable for both male and female adults, but strongly influenced by various combinations of fungal isolate, exposure stage, and adult sex.

## Discussion

This study describes a new *B. bassiana* isolate, which was recovered from Ae. albopictus L4 larvae, collected from artificial containers in Manhattan, KS, USA in August 2018. Using a simple laboratory-based experimental design, this study reports the impact of the MHK isolate, as well as the commercially available GHA isolate, on multiple life history parameters of *An. gambiae* and *Ae. albopictus* mosquitoes. We acknowledge that, in the field, efficacy of fungal biocontrol agents for mosquito control are influenced by many factors, including abiotic and biotic conditions, types of formulations, and deployment, as well as the broader ecological context of disease transmission (Clark et al. 1968, Rangel et al. 2004, Bukhari et al. 2011, Fang and Leger 2012). This study solely aimed to determine the initial potential of the *B. bassiana* MHK isolate as a source reduction tool for the control of *An. gambiae* and *Ae. albopictus*. Our results show that *B. bassiana* isolates MHK and GHA reduced larval survival in both mosquito species. However, *An. gambiae* larvae were more susceptible to infection compared to *Ae. albopictus* larvae. We reveal that irrespective of mosquito species, larval exposure to *B. bassiana* delayed development to pupae and adults and further reduced the survival of emerged male and female mosquitoes.

Previous studies have reported isolate-specific virulence of entomopathogenic fungi, including *B. bassiana*, towards *Ae. albopictus* mosquito larvae (Chitra and Phil 2011), larvae of the cabbage moths (Batcho et al. 2018), and adult house flies (White et al. 2021). Over study did not reveal such isolate-specific virulence of *B. bassiana* MHK on neither *An. gambiae* nor *Ae. albopictus* larvae. Indeed, the MHK isolate was as efficient in reducing larval survival and impacting mosquito development as the GHA isolate. Similar results have been reported in studies assessing the entomopathogenic activity of local *B. bassiana* isolates towards several insects, including mealybug adults and red palm weevils, as compared to commercially available isolates (Sutanto et al. 2021, Mathulwe et al. 2022). While we acknowledge that statistical evaluation of efficacy between the two fungal isolates is not possible in our study, as larval exposures were conducted using different mosquito generations, our study confirms that the entomopathogenicity of *B. bassiana* on mosquito larvae is maintained in the MHK isolate. The larvicidal activity of MHK isolate is consistent with three previous studies reporting above 90% mortality in *An. gambiae* early and late instar larvae (Bukhari et al. 2010, Prasad and Veerwal 2012), and up to 46% mortality in *Ae. albopictus* larvae (JianCong et al. 2013) exposed to various *B. bassiana* isolates.

*B. bassiana* MHK and GHA isolates reduced larval survival of *An. gambiae* and *Ae. albopictus* in a dose-dependent manner between the low and the two highest fungal doses. This result was expected and is in agreement with previous reports for *An. gambiae* and *Anopheles stephensi* Liston (Diptera: Culicidae) larvae exposed to entomopathogenic fungi, including *B. bassiana, Isaria spp*., and *M. anisopliae* (Bukhari et al. 2010, Prasad and Veerwal 2010, Ramirez et al. 2018). However, our data show that difference in larval mortality remained minimal between the medium and high fungal doses. While we cannot exclude that this observation is explained by a narrow concentration gradient of *B. bassiana’s* larvicidal activity, it may also be explained by the type of conidial formulation used in this study that solely relied on water. The hydrophobic fungal conidia of *B. bassiana*, when applied at high concentrations, tend to clump and sink to the bottom of the mosquito breeding habitat (Bukhari et al. 2010). The addition of synthetic oil as a carrier in *B. bassiana* and *M. anisopliae* conidial formulations increases the ability of the fungal conidia to spread on the water surface and increases larval mortality in *An. gambiae* mosquito larvae (Bukhari et al., 2011). While the impact of different formulations on the efficacy of B. bassiana MHK was beyond the scope of this study, future study designs should assess whether synthetic oils can further enhance the larvicidal activity of the *B. bassiana* MHK isolate towards mosquito larvae.

Our data show that the larvicidal activity of *B. bassiana* was much stronger against *An. gambiae* than *Ae. albopictus*. This result agrees with previous findings that compared directly the larvicidal activity of *B. bassiana* against several mosquito species. *B. bassiana* was more effective in killing larvae of several *Anopheles* and *Culex* species than killing *Aedes aegypti* (L.) (Diptera: Culicidae) and/or *Aedes sierrensis* (Ludlow) (Diptera: Culicidae) larvae (Clark et al. 1968, Geetha and Balaraman 1999). The lower mortality observed in *Aedes* larvae may be attributed to a shorter larval stage and the reduced ability of *B. bassiana* to penetrate the perispiracular lobes of *Aedes* larvae compared to *Anopheles* and *Culex* (Clark et al. 1968, Geetha and Balaraman 1999). In addition, previous studies have shown that *B. bassiana* blastospores have a higher larvicidal effect against *Ae. aegypti* larvae as compared to conidiospores (Miranpuri and Khachatourians 1990, 1991). Thus, future studies could test *B. bassiana* MHK blastospores in infection bioassays and utilize histological assays to determine whether similar infection patterns are observed in *An. gambiae* and *Ae. albopictus* mosquito larvae.

Previous studies demonstrated that the efficacy of larvicidal activity of entomopathogenic fungi is influenced by the age of mosquito larvae at the time of exposure. In general, larval mortality was higher when early instar as compared to late instar larvae of *An. stephensi*, *Ae. aegypti*, *Cx. quinquefasciatus, or Culex tritaeniorhynchus* Giles (Diptera: Culicidae) were exposed to *B. bassiana* or *M. anisopliae* (Sandhu et al. 1993, Geetha and Balaraman 1999, Bukhari et al. 2010, Prasad and Veerwal 2012). Increased susceptibility of younger larvae may be explained by overall longer exposure to fungal conidia and presence of fungal conidia during more molts, where thinner cuticle is exposed (Bukhari et al. 2010, Sandhu et al. 1993). Our data showed a similar pattern for *Ae. albopictus* larval mortality after exposure to either the MHK, or the GHA isolate of *B. bassiana*. This pattern was reversed for *An. gambiae*, where fungal exposure at the L3 stage killed more mosquito larvae as compared to exposure at the L2 stage. The increased efficacy of larvicidal activity with increased larval age was subtle, as the vast majority of *An. gambiae* larvae died irrespective of exposure age. A previous study on the impact of *B. bassiana* on *An. gambiae* larval survival also failed to detect increased larvicidal activity when younger larval stages were exposed (Bukhari et al. 2010). Currently, the literature on larvicidal effects of entomopathogenic fungi on *An. gambiae* is limited, and future studies are required to determine whether later stages of *An. gambiae* are in general more susceptible to entomopathogenic fungi.

In addition to larvicidal activity, fungal entomopathogens have the potential to delay insect development. Indeed, our study shows that larval exposure to *B. bassiana* MHK delays the development of *Ae. albopictus* and *An. gambiae* larvae to adults by up to two and four days, respectively. These results corroborate studies reporting increased developmental time to pupae and adults in *Ae. albopictus* and *Cx. pipiens* larvae exposed to various entomopathogenic fungi, including *B. bassiana* and *M. ansiopliae* (Shoukat et al. 2016, 2018, 2020). Delay in larval development has also been reported in the larvae of several moth species, including corn earworm, greenish silk-moth, and gypsy moth exposed to *B. bassiana, Isaria fumosorosea, M. anisopliae*, and *Nosema necatrix* (Mitchell and Cali 1994, Henn and Solter 2000, Hussain et al. 2009). The observed delay in development in moth larvae has been attributed to physiological changes and reduced larval feeding induced by fungal infection (Henn and Solter 2000, Hussain et al. 2009). While the underlying cause of developmental delay in mosquitoes has yet to be determined, this delay extends the mosquito population doubling time and thus further decreases population growth.

Furthermore, our data demonstrate that larval exposure to *B. bassiana* MHK reduced pupal and adult survival in both mosquito species, and thus negatively impacts the life history of *An. gambiae* and *Ae. albopictus* beyond the larval stage. Our results agree with previous studies that report fungus-induced mortality in the pupae of *An. gambiae* and *An. stephensi* mosquitoes exposed to *B. bassiana* as larvae (Sandhu et al. 1993, Pereira et al. 2009, Bukhari et al. 2010, 2011, Prasad and Veerwal 2012, Vogels et al. 2014, Veerwal et al. 2022). However, to our knowledge this is the first report of increased adult mortality after exposure of mosquito larvae to *B. bassiana*. The underlying cause of this adult mortality is currently unknown, but may be explained by impaired larval physiology reducing adult survival or an active fungal infection in mosquito adults. The latter is partially supported by a report of *B. bassiana*-infected *An. gambiae* and *An. stephensi* pupae and adults following larval exposure (Bukhari et al. 2010). The source of this infection is unknown, as the experimental design of Bukhari et al., as well as ours, is unable to distinguish between trans-stadial infection and re-acquisition of the infection after eclosion from the larval environment. Regardless of the underlying cause, the use of *B. bassiana* as a larvicide reduces mosquito survival beyond the larval stage.

In conclusion, this study provides a first report on the isolation and characterization of a mosquito-associated *B. bassiana* isolate, MHK, with entomopathogenic activity against two mosquito species of public health concern. Using a straight-forward laboratory-based experimental design, our data suggest that the *B. bassiana* MHK isolate is a potential larvicide for *An. gambiae*. Exposure of *An. gambiae* larvae to *B. bassiana* MHK did not only reduce larval, pupal, and adult mosquito survival, but also significantly increased development time, parameters that combined have the potential to strongly impair mosquito population growth. In addition, our results suggest that further evaluation of the potential use of the MHK isolate for *Ae. albopictus* control is warranted, given its substantial negative impact on pupal survival.

## Acknowledgements

We acknowledge that larval collections and our research was conducted in Manhattan, KS, on land historically home to many Native nations, including the Kaw, Osage, and Pawnee, among others. We further acknowledge that Kansas State’s history as the first land grant university in the United States rests on the dispossession of Indigenous peoples and nations from their lands. This study was supported by funding from National Institutes of Health grant number R01AI140760, USDA-ARS specific cooperative agreement 58-5430-4-022, and USDA National Institute of Food and Agriculture Hatch project 1021223. This is contribution no. 22-313-J from the Kansas Agricultural Experiment Station. The contents of this article are solely the responsibility of the authors and do not necessarily represent the official views of the funding agencies.

