## Supplementary Fig. S1 for "The potential of the *Beauveria bassiana* MHK isolate for mosquito larval control"

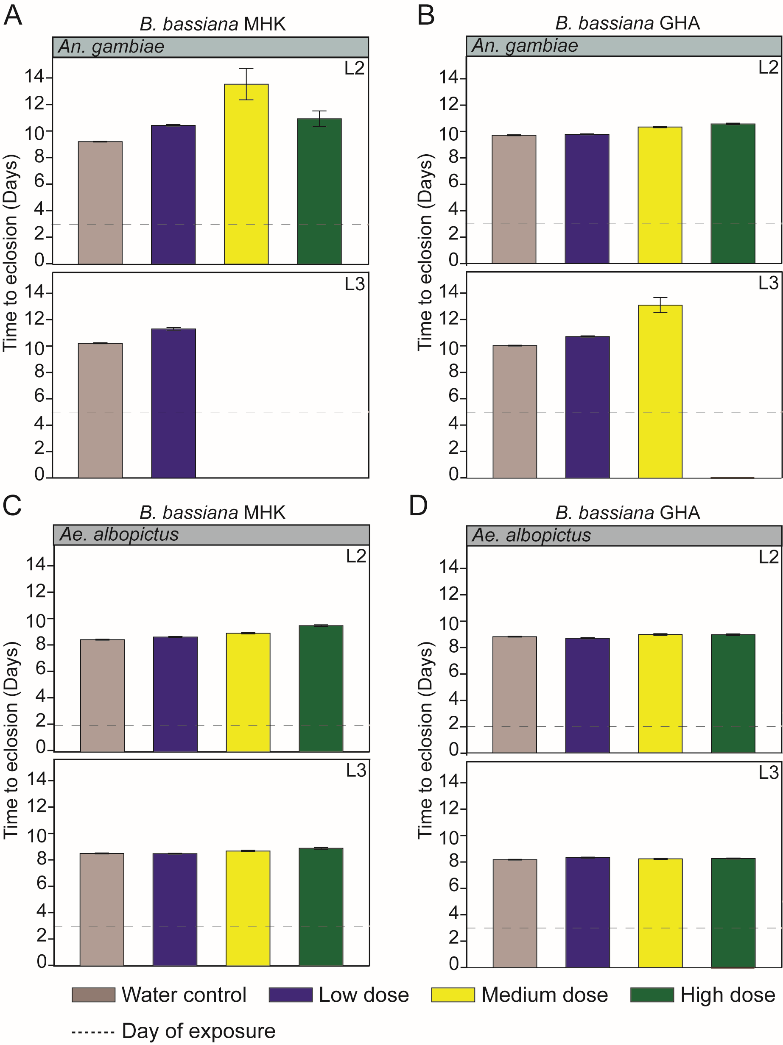

**Supplementary Fig. 1.** Mean time to eclosion in days (Eclosion being Day 0) of L2 and L3 larvae exposed to *B. bassiana*. Larval exposure stage to *B. bassiana* is indicated at the top right of each bar graph. Dotted line indicates the day each larval stage was exposed to *B. bassiana*. The error bar indicates the standard error of mean time to eclosion. (**A**) Mean time to eclosion of *An. gambiae* larvae exposed to *B. bassiana* MHK; (**B**) Mean time to eclosion of *An. gambiae* larvae exposed to *B. bassiana* GHA; (**C**) Mean time to eclosion of *Ae. albopictus* larvae exposed to *B. bassiana* MHK; (**D**) Mean time to eclosion of *Ae. albopictus* larvae exposed to *B. bassiana* GHA.
