## Supplementary Table S5 for "The potential of the *Beauveria bassiana* MHK isolate for mosquito larval control"

**Supplementary Table 5. Time to eclosion in days (Eclosion being Day 0) for *An. gambiae* and *Ae. albopictus* L2 and L3 larvae exposed to *B. bassiana* MHK and GHA.**

|  | | | | | **Difference (P-value)** | | |
| --- | --- | --- | --- | --- | --- | --- | --- |
| ***B. bassiana* strain** | **Mosquito species** | **Larval stage** | **Fungal dose** | **LSM*+/-SEM** (Days)** | **Control** | **Low dose** | **Medium dose** |
| MHK | *An. gambiae* | L2 | Control | 9.29 +/- 0.03 | - | - | - |
|  |  |  | Low | 10.5 +/- 0.06 | 1.23 (<.001) | - | - |
|  |  |  | Medium | 13.6 +/- 1.19 | 4.36 (0.067) | 3.12 (0.119) |  |
|  |  |  | High | 11.0 +/- 0.59 | 1.74 (0.003) | 0.510 (0.392) | -2.62 (0.140) |
|  |  | L3 | Control | 10.2 +/- 0.04 | - | - | - |
|  |  |  | Low | 11.3 +/- 0.10 | 1.09 (<.001) | - | - |
|  | *Ae. albopictus* | L2 | Control | 8.50 +/- 0.04 | - | - | - |
|  |  |  | Low | 8.70 +/- 0.04 | 0.200 (<.001) | - | - |
|  |  |  | Medium | 9.00 +/- 0.05 | 0.490 (<.001) | 0.290 (<.001) | - |
|  |  |  | High | 9.57 +/- 0.07 | 1.06 (<.001) | 0.860 (<.001) | 0.570 (<.001) |
|  |  | L3 | Control | 8.49 +/- 0.04 | - | - | - |
|  |  |  | Low | 8.48 +/- 0.04 | -0.010 (0.779) | - | - |
|  |  |  | Medium | 8.68 +/- 0.05 | 0.190 (0.003) | 0.200 (0.001) | - |
|  |  |  | High | 8.88 +/- 0.06 | 0.390 (<.001) | 0.400 (<.001) | 0.200 (0.014) |
| GHA | *An. gambiae* | L2 | Control | 9.68 +/- 0.04 | - | - | - |
|  |  |  | Low | 9.76 +/- 0.04 | 0.080 (0.181) | - | - |
|  |  |  | Medium | 10.3 +/- 0.05 | 0.620 (<.001) | 0.550 (<.001) | - |
|  |  |  | High | 10.5 +/- 0.06 | 0.860 (<.001) | 0.780 (<.001) | 0.240 (0.003) |
|  |  | L3 | Control | 10.0 +/- 0.04 | - | - | - |
|  |  |  | Low | 10.7 +/- 0.05 | 0.680 (<.001) | - | - |
|  |  |  | Medium | 13.1 +/- 0.58 | 3.07 (0.033) | 2.38 (0.052) | - |
|  | *Ae. albopictus* | L2 | Control | 8.80 +/- 0.04 | - | - | - |
|  |  |  | Low | 8.68 +/- 0.05 | -0.120 (0.065) | - | - |
|  |  |  | Medium | 8.97 +/- 0.07 | 0.170 (0.031) | 0.280 (0.001) | - |
|  |  |  | High | 8.95 +/- 0.07 | 0.150 (0.051) | 0.270 (0.002) | -0.020 (0.869) |
|  |  | L3 | Control | 8.18 +/- 0.03 | - | - | - |
|  |  |  | Low | 8.34 +/- 0.05 | 0.150 (0.006) | - | - |
|  |  |  | Medium | 8.25 +/- 0.04 | 0.070 (0.198) | -0.090 (0.144) | - |
|  |  |  | High | 8.27 +/- 0.04 | 0.090 (0.097) | -0.070 (0.271) | 0.020 (0.713) |

LSM*: Least squares mean; SEM**: Standard error of mean.
